# Metabolic and Hormonal Alterations Induced by Roux-en-Y Gastric Bypass

**DOI:** 10.1101/230201

**Authors:** Loqmane Seridi, Gregory C. Leo, G. Lynis Dohm, Walter J. Pories, James Lenhard

## Abstract

Roux-en-Y gastric bypass (RYGB) is an effective way to lose weight and reverse type 2 diabetes. We profiled the metabolome of 18 obese patients (nine euglycemic and nine diabetics) that underwent RYGB surgery and seven lean subjects. Plasma samples from the obese patients were collected before the surgery and one week and three months after the surgery. We analyzed the metabolome in association to five hormones (Adiponectin, Insulin, Ghrelin, Leptin, and Resistin), four peptide hormones (GIP, Glucagon, GLP1, and PYY), and two cytokines (IL-6 and TNF). PCA showed samples cluster by surgery time and many microbially driven metabolites (indoles in particular) correlated with the three months after the surgery. Network analysis of metabolites revealed a connection between carbohydrate (mannosamine and glucosamine) and glyoxylate and confirms glyoxylate association to diabetes. Only leptin and IL-6 had a significant association with the measured metabolites. Leptin decreased immediately after RYGB (before significant weight loss), whereas IL-6 showed no consistent response to RYGB. Moreover, leptin associated with tryptophan in support of the possible role of leptin in the regulation of serotonin synthesis pathways in the gut. These results suggest a potential link between gastric leptin and microbial-derived metabolites in the context of obesity and diabetes.

## Introduction

Obesity and type 2 diabetes (T2D) are among the top preventable causes of death worldwide (obesity kills ~3 million yearly) [1]. The number of obese and diabetic patients doubled in less than three decades [2,3]. The rapid rise of obesity and T2D reflects a complex interaction between the fast-changing environment (e.g., lifestyle) and the slow adapting biology (genetics). This complexity has led to diverse preventive and treatment approaches to address both the environmental (diet and exercise) and biological (medications and surgeries) aspects of the diseases; however, the effects of the current approaches are limited.

Roux-en-Y gastric bypass (RYGB) is a surgical procedure that creates a small pouch from the stomach while bypassing the main portion of the stomach and most of the duodenum. The small pouch is connected to the jejunum forming a “Y” with the bypassed stomach and duodenum components of the digestive tract. RYGB achieves remarkable results in addressing obesity and T2D - it persistently decreases weight and hyperglycemia in most patients [4,5]. Moreover, targeted metabolomic analyses showed that RYGB reduces circulating metabolites implicated in obesity and insulin resistance such as branch chain amino [6,7] and ceramides [8,9]. Untargeted metabolomic analyses confirm and unveiled alterations of essential metabolites by RYGB [10,11].

We conducted an untargeted metabolomics analysis of plasma from obese and obese diabetic patients in comparison with plasma from healthy lean women. Also, we studied the metabolic alterations by RYGB in association to distinct clinical features plus nine protein and peptide hormones and two cytokines.

## Results

### RYGB alters the metabolome profiles of obese patients independently of disease state

We studied a cohort of 27 Caucasian women [12]: 18 obese (nine diabetics and nine nondiabetics; BMI>35 kg/m^2^) and nine lean subjects (BMI<25 kg/m^2^). We dismissed two lean subjects because of incomplete data. Hereafter, we use “obese’ to refer to both obese diabetic and obese non-diabetic patients. Obese patients underwent RYGB surgery causing weight loss and improvement of pre-diabetic/diabetic symptoms— rapid normalization of circulating insulin x(within one week) and glucose (three months) [12] (Supplementary Fig. S1).

To understand the effects of RYGB on metabolic activities, we profiled the metabolome of the obese subjects one week before the surgery and one week and three months after the surgery; and of the lean subjects as a control. We measured 223 (including unknowns) and focused on 148 known metabolites (10 carbohydrates, 32 amino acids, and 80 complex lipids).

Principal component analysis (PCA) revealed a separation between obese and lean subjects (Fig. 1a). Moreover, samples from obese subjects cluster by surgical stage (pre-surgery, one week and three months after surgery) rather than disease state (diabetic vs. non-diabetic). Indeed, this clustering is more prominent when we excluded samples of lean subjects (Fig. 1b). Finally, we validated these observations using partial least square discriminate analysis (PLS-DA) (Supplementary Fig. S2).

**Figure 1:**
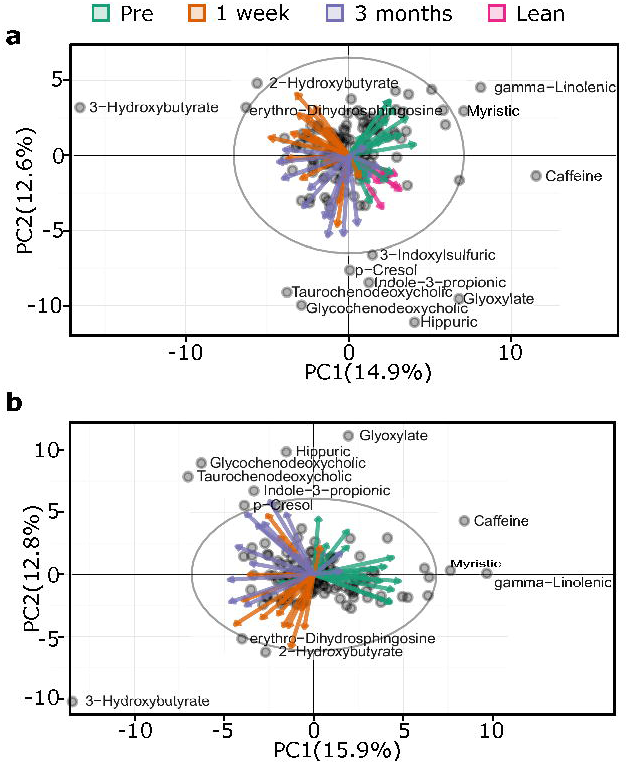
Principal component analysis and partial least square discriminant analysis of metabolomics profiles (148 metabolites and 25 patients). Projection of metabolic data a) PCA using all subjects, it shows patients correlate with surgical condition b) same as (a) but excluding lean; eclipses represent 95% confidence interval. For visualization, only scattered metabolites are labeled.

In sum, PCA indicates that the metabolome of obese patients was different from that of lean subjects. RYGB affected the metabolome of obese diabetic and obese non-diabetic similarly. Moreover, each surgical stage correlates with specific metabolites.

### Network analysis unveil a connection between glyoxylate and hexosamines

To understand the association between the measured metabolites, we constructed an information-theory-based metabolite network (Figure 2). The network is structured into modules of strongly interconnected lipids, amino acids, and carbohydrates. Interestingly, one module connects hexosamines (mannosamine and glucosamine) to members of the glyoxylate cycle (glyoxylate, oxalate, and succinate and fumarate). We inspected the response of these metabolites to RYGB. As expected, mannosamine and glucosamine dropped in response to RYGB in diabetic patients (P=0.02). Oxalate, succinate, and fumarate did not respond to RYGB and did not exhibit any significant difference between diabetic and none-diabetic obese patients (P=0.27, 0.32, 0.93, respectively). Glyoxylate showed a trending decrease in response to RYGB (though not statistically significant); however, glyoxylate was higher in diabetic compared to none-diabetic obese patients (P=0.008).

**Figure 2:**
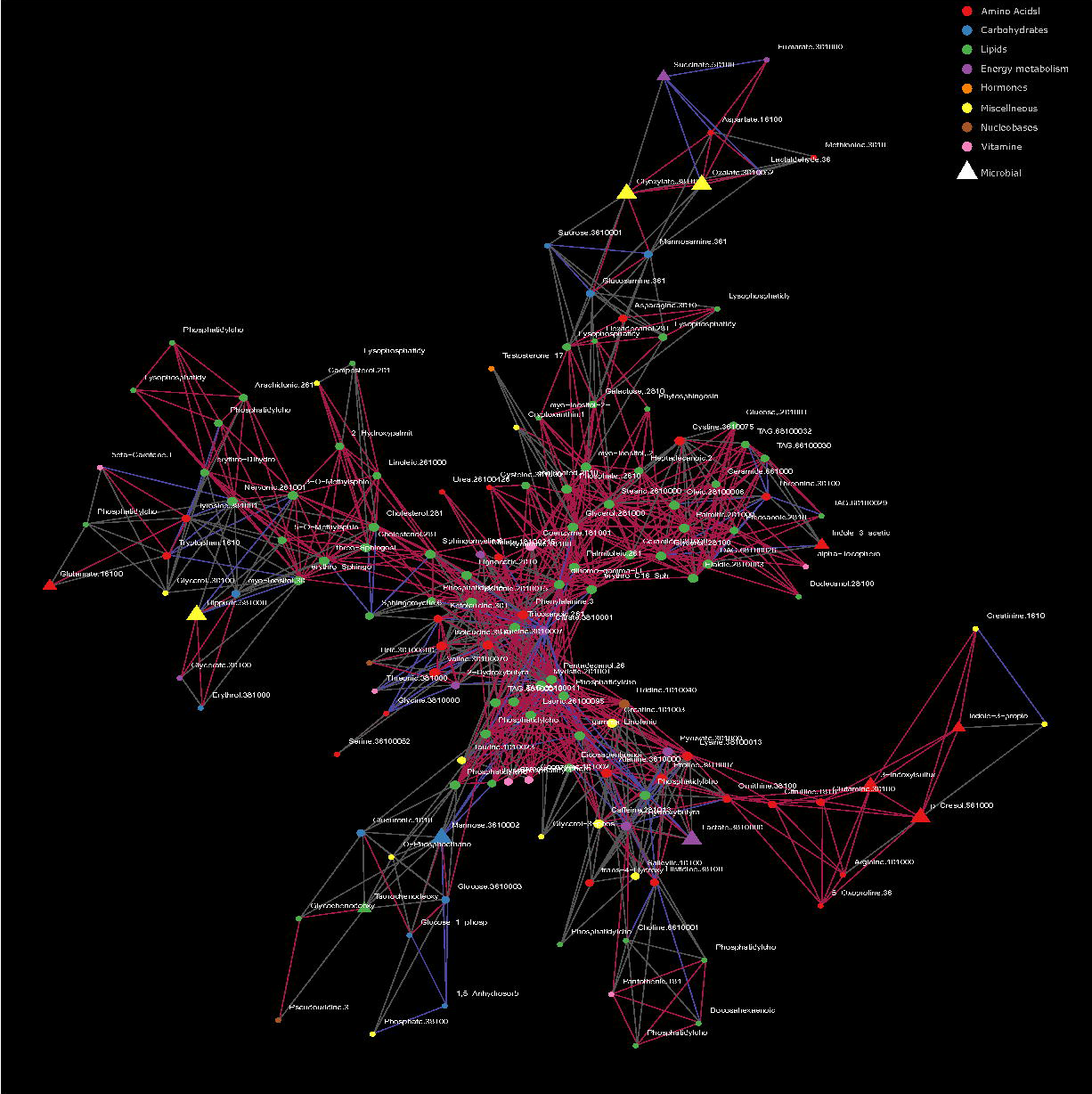
Metabolic network. Association network between metabolites. Red edges indicate positive correlation (Spearman>0.2); blue indicates negative correlation (Spearman<-0.2); and gray are indicative of possible association but no significant correlation. Node size correlates with node degree.

### Metabolites correlate with Leptin and IL-6

In addition to the metabolic profiling and network analysis, we evaluated the metabolic alterations associated with the following parameters: fasting plasma glucose (FPG); C-peptide; free fatty acid (FFA); five hormones (Adiponectin, Insulin, Ghrelin, Leptin, and Resistin); four peptide hormones (GIP, Glucagon, GLP1, and PYY); and two cytokines (IL-6 and TNF). For simplicity, we will refer to these measurements as “clinical features”.

To investigate the associations between the metabolites and the clinical features, we conducted orthogonal partial least square (OPLS) analysis. 5 out of 14 clinical features (FPG, BMI, weight, leptin, and IL-6) associated with at least one metabolite and 70 metabolites associated with at least one clinical feature (VIP>=1.5, Q^2^(cum)>=0.4 and R^2^Y>=0.5; Fig. 3). We validated the robustness of the OPLS models by the row permutation test [13] (Supplementary Fig. S3). Also, we checked whether it could capture expected associations. Indeed, FPG linked with carbohydrates (mannosamine, glucosamine, and glucose) and 1,5 anhydrosorbitol (1,5AG); the latter is consistent with depletion of 1,5AG at hyperglycemia [14]. Furthermore, we confirmed associations of FPG, weight, and BMI to plasma amino acids [15–17]; and BMI and body weight to complex fatty acids and lipids [18].

**Figure 3:**
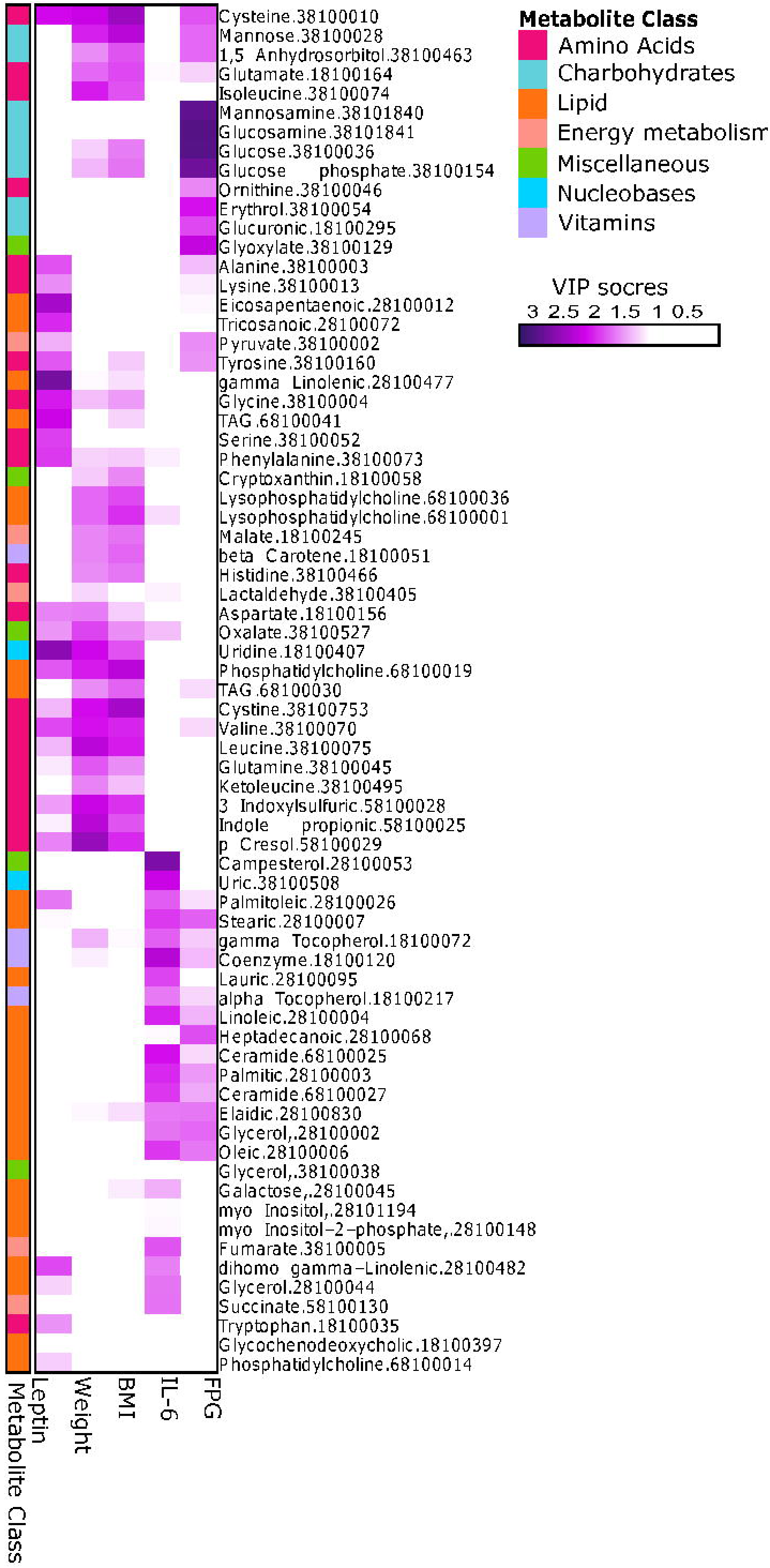
Metabolites associations to clinical features. The heatmap shows “clinical features” (columns) association to metabolite (rows). The heatmap shows metabolites/clinical feature that has at least one association (VIP>1.5). Metabolites are ordered based on hierarchical clustering. For clarity, only part of metabolite name and unique IDs are shown. The full description of a metabolite can be queried by the ID in Supplemental Table S1.

Figure 3 shows that leptin lacks association with carbohydrates, but has a strong association with amino acids (e.g., leucine and tryptophan), complex fatty acids (e.g., gamma-linolenic and eicosapentaenoic acids) and lipids. Also, IL-6 is independent of carbohydrates and amino acids but correlates with various complex fatty acids and lipids (e.g., ceramide and palmitic acid).

### Leptin and IL-6 response to RYGB

Leptin is a hormone secreted by adipose tissues and by gastric mucosa[19]. It regulates food intake [20] and enhances energy expenditure [21]. Leptin is a hormone linked to fat stores and diseases such as anorexia nervosa, obesity, and Alzheimer’s [22,23]. Before the surgery, we found fasting plasma leptin higher in obese patients compared to obese diabetic patients (P=0.004) and compared to lean (P=3.35e-5) subjects. Also, leptin in obese diabetic patients was comparable to that of leans subjects (P=0.12). Leptin declined immediately (1 week) after RYGB surgery (nondiabetic, P=6.32e-6; diabetic, P=0.005) and fell below the level in lean subjects after three months (nondiabetic, P=0.06; diabetic, P=0.003).

IL-6 is a cytokine secreted by immune, fat and muscle cells. IL-6 responds to various stresses [24] and acts as both pro and anti-inflammatory agent [25]. Circulating IL-6 correlates with insulin resistance [26]. As expected [27], IL-6 is elevated in obese compared to lean subjects and the trend remains higher after three months (nondiabetic, P_pre_=0.04, P_3months_=0.09; diabetic, P_pre_=0.02, P_3months_=0.02). IL-6 does not change in respond to RYGB (non-diabetic, P_1week/pre_=0.77, P_3months/pre_=0.59; diabetic, P_1week/pre_=0.51, P_3months/pre_=0.89).

## Discussion

We profiled the metabolome of obese diabetic and obese non-diabetic patients before RYGB, and one week and three months after RYGB, and of lean subjects. The metabolome profile of obese is distinct from that of lean patients. RYGB affected the metabolome of both obese groups similarly, confirming previous report [11]. Each surgical stage correlated with specific metabolites. For instance, 3-hydroxybutyrate correlates with the one-week post-surgery, possibly because of the beta-oxidation of fats due to the change in diet; i.e., from Optifast 800 (Novartis Nutrition Group, Vevey, Switzerland) to Bariatric Advantage Meal Replacement (Bariatric Advantage, Irvine, CA) after surgery [12]. The 2-hydroxybutyrate may reflect production of glutathione due to the oxidative stress arising from the surgical procedure. The erythro-dihydrosphingosine or sphinganine is consistent with its role in regulating CD-95 mediated apoptosis of T-cells [28], possibly, contributing to post surgery recovery. Metabolites of microbial origin (indole-3-propionic acid, 3-indolesulfuric acid, hippuric acid and glycochenodeoxycholate acid) [29–31] correlated with the three months post-surgery; which suggests a change in the gut microbial population. Indeed, metagenomics sequencing before and after three months RYGB surgery showed a reduction of *Firmicutes* and *Bacteroidetes* and an increase of *Proteobacteria* and *Verrucomicrobia* [32]. Moreover, in *in-vitro* models, increasing indole-3-propionic acid reduced inflammation and intestinal permeability and altered the glucose/fructose transporter GLUT5 mRNA transcription [33].

Our network analysis showed an association between carbohydrates (mannosamine and glucosamine) and glyoxylate. Mannosamine and glucosamine are substrates for O-linked protein glycosylation and precursors for sialic acids which are important in N-linked protein glycosylation. O-linked glycosylation and the hexosamine biosynthetic pathway have been implicated in causing insulin resistance [34]. The hexosamines were elevated in diabetic patients and decreased in response to RYGB. We speculate that the alteration of gut microbes may, in part, contribute to the decrease in circulating hexosamines through altered breakdown of mucin within the intestine. The glyoxylate was elevated in diabetic patients which supports the suggested glyoxylate link to [35,36]. Glyoxylate is produced through the glyoxylate cycle which is an anabolic pathway that converts acetyl-CoA to glyoxylate and succinate in plants, bacteria, and invertebrates [37]. The succinate is used to produce cellular carbohydrates, and the glyoxylate is converted to malate to complete the abbreviated metabolic cycle. Change in glyoxylate levels could reflect alterations of the microbiome; indeed, *Proteobacteria* is higher in the microbiome of diabetic patients compared to non-diabetic patients [38]. Although *Proteobacteria* increased after RYGB, this increase is reported for non-diabetic patients; and the effects of RYGB on microbiome of diabetic patients is required. The gut microbial origin of glyoxylate is consistent with nutrients-depleted environment expected in the lower gut. The microbial glyoxylate shunt is expected to be active in such environment, and the fuel for the shunt could be acetate produced by commensal gut microbes. In a long-term study of the effects of bariatric surgery, gut microbiota composition for *Escherichia*, *Klebsiella*, and *Salmonella*, among other species, increased in women who had RYGB versus obese women [39]. This same study showed that glyoxylate and dicarboxylate metabolism was enriched in vertical banded gastroplasty patients versus obese women but not for RYGB patients. The ability of any of these bacteria to degrade intestinal mucin was not discussed. Alternatively, glyoxylate could arise from glycine and hydroxyproline catabolism in the liver [40], although, our network analysis did not show a correlation of glyoxylate with glycine or hydroxyproline while a correlation with oxalate was observed. The network analysis allows for possible biochemical interpretation, but as previously mentioned glyoxylate, succinate and oxalate measured in the plasma did not respond to RYGB; although glyoxylate was higher in diabetic compared to non-diabetic patients.

Leptin declined immediately after RYGB and fell below the level in lean subjects after three months. Indeed, other studies reported similar decline one [41], two [42] and three [43] weeks after the RYGB. The decrease in stomach secreted leptin due to RYGB alteration of the stomach’s physiology is speculated to cause this immediate response [42]. Gastric epithelial cells adapted to high-fat diet by increasing gastric leptin secretion in fasting mice [44]. Therefore, it is possible that adaptation to a new diet after RYGB may cause a decrease in gastric leptin. Also, gastric leptin (both protein [45] and mRNA) is higher in obese compared to lean subjects, and changes in gastric leptin precede changes in plasma leptin [44]. Circulating leptin, in part, originates from gastric leptin, most notably after food intake [19]. In sum, leptin drops in response to RYGB before any significant weight loss. This response could be due to decreases of gastric leptin secretion caused by the changes in the physiology of the stomach [42] or the changes in diet or both.

OPLS analysis shows that leptin correlated with several amino acids, including branched-chain amino acids. This is consistent with the observation that leucine-supplemented diet given to ob/ob mice for two weeks increased plasma leptin [46]. In rats, leucine-deficient diet reduced leptin response to meals (diet deficient in other amino acids did not alter leptin response to meals) [47]. Leucine enhances leptin sensitivity in rats on a high-fat diet [48] and regulates leptin translation (rather than transcription) in adipose tissues in rats. Moreover, plasma total cysteine positively correlates with leptin in a Hispanic cohort (the correlation is partially independent of fat mass) [49] and alanine stimulated leptin expression in rats [50].

The leptin association to tryptophan was unexpected. For example, tryptophan composition in diet yielded inconsistent leptin responses in pigs [51]. Recently, gastric leptin was proposed to regulate serotonin synthesis pathways through a mechanism involving increased expression of tryptophan hydroxylase-1 (TPH1; a rate limiting enzyme that converts tryptophan to serotonin) in the gut of obese subjects (similar to gastric leptin). Also, oral administration of leptin increased expression TPH1 in the intestine of leptin-deficient mice [44]. One hypothesis is that the association between circulating leptin and tryptophan may be partially explained by gastric leptin regulation of the tryptophan-serotonin pathways in the gut.

Another possible explanation of the leptin-tryptophan association could be through the aryl hydrocarbon receptor (AHR). AHR has been implicated in mice to play a significant role in obesity [52]. Tryptophan-derived indole metabolites (kynurenine, tryptamine, indole-3-acetic acid, indole-3-aldehyde and indole-3-acetaldehyde) have been shown to modulate AHR [53]. Several tryptophan-derived indole metabolites were observed in this study. We observed kynurenic acid decreased after RYGB which is in agreement with suggested kynurenine acid association with higher BMI [54]. Furthermore, Figure 3 showed associations between leptin, weight, and BMI and 3-indoxylsulfate and indole-3-propionic acid. Interestingly, removal of gut microbes by antibiotic treatment increases tryptophan and reduces tryptophan-related metabolites (i.e., serotonin and indole derivatives) in circulation and decreases weight gain. Xu et al. reported that, under a high-fat diet, AHR-deficient mice produced less leptin than wild-type mice. They suggested that the difference is due to the reduced amounts of epididymal white adipose tissue; however, the effects on the gut microbiome were not investigated [55]. Moreover, mice exposed to 2, 3, 7, 8-tetrachlorodibenzofuran, an AHR ligand, stimulated lipogenesis [56]. We report a decrease in the AHR ligand, kynurenic acid, after RYGB surgery. Shin et al. in exploring the role of NRF2 (NF-E2 p45-related factor 2) pathway reported that it activated the AHR signaling cascade resulting in an inhibition of adipogenesis [57]. This negative regulation by AHR of adipogenesis, the authors reported was consistent with previous literature for one of the roles of AHR. Summarizing the present work, these data support the suggested correlation between leptin, body weight and the microbiome [58–61], however, whether leptin directly affects the microbiome and indole-mediated AHR signaling needs further investigation.

In this study, IL-6 did not respond to RYGB. Previous studies were contradictory regarding IL-6 response to RYGB. After RYGB: IL-6 increased within the first week and remained high at three months and decreased after one year [41]; IL-6 decreased after six months [27] and one year [62]; and IL-6 did not change within three weeks [63], one month [62], three months, six months [63] and one years [64]. These inconsistent results may reflect complex inflammatory patterns among obese and diabetic patients in response to uncontrolled environmental influences. Our observation that IL-6 correlates with various lipids (e.g., ceramide and palmitic acid), but not carbohydrates or amino acids, is consistent with the lipid signaling and IL6-mediated stress responses crosstalk in metabolic diseases [57].

## Conclusion

We have presented our analysis of untargeted metabolomics of plasma from obese and obese diabetic female patients in comparison with plasma from healthy lean women and associated the metabolic alterations by RYGB to distinct clinical features. What stands out in our results is the strong connection with the microbial metabolites at three months post-surgery and the connection between leptin and tryptophan and indole-related metabolites. The microbiome connection is consistent with metagenomics analyses of RYGB patients before and after surgery, and the leptin connection opens a new avenue of consideration.

## Methods

### Cohort, RYGB surgery procedure, and clinical data measurements

The cohort and the procedure of RYGB surgery are described in [12]. All clinical features were measured as part of that study, but we are reporting them for the first time.

### Metabolomic profiling

The metabolome was profiled by Metanomics Health (Berlin, Germany) using their broad 4 phases profiling approach of gas chromatography-mass spectrometry (GC-MS) and liquid chromatography-mass spectrometry (LC-MS/MS) [65,66].

### Statistical analysis

R 3.2.4 was used for statistical analysis. “ropls” package [67] was used for PLS-DA and OPLS analysis. Values outside the 1.5 interquartile range were discarded as outliers. Figures were generated using ggplot2, and pheatmap of R. Two-sided t-test was used for all comparisons.

### Network analysis

Scores based on information variation over minimum spanning tree was computed between metabolites. Insignificant links were removed after calculating expected scores based on 1000 stochastic simulations. The network was visualized using ggnet2.

## Author contribution

JL conceived the overall study. LS and GL analyzed the data. JL, GL, LS interpreted the data. LS, GL, and JL wrote the manuscript. GLD and WJP designed the experiment, assisted in the collection of the data and reviewed the manuscript.

## Competing interests

JL, GL, and LS are employees of Janssen Research & Development, LLC and have no other conflict of interest to disclose.

---

**Supplemental Figure S1: Decrease in weight, fasting insulin and fasting glucose after RYGB**. a) Barplot shows the number of subjects participated in the study. b) Boxplot shows the age distribution of the participants; diabetic participants are significantly older than the others c) Boxplot shows a significant weight loss of patients after RYGB. d) Same as (c) but for BMI. e) Barplot shows a significant drop of fasting insulin levels within the first week. f) shows a significant decline in fasting glucose levels within the first week of RYGB. We defined outlier points by the values outside the 1.5 interquartile range. We omitted outliers from the analysis. This figure is a representation of previously published data [12].

**Supplemental Figure S2: Partial least square discriminant analysis of metabolomics profiles (148 metabolites and 25 patients).** Projection of metabolic data a) PLSDA using all subjects, it shows a correlation between samples of the same surgical condition b) same as (a) but excluding lean subjects; eclipses represent 95% confidence interval.

**Supplemental Figure S3: Permutation tests for orthogonal partial least square analysis.** Boxplots show the distribution of Q^2^ and R^2^ for 1000 models. Each model generated after the rows of the metabolic data were randomly permuted.

**Table S1: Metabolome data.** The list of measured metabolites.

